# DANSE: a pipeline for dynamic modelling of time-series multi-omics data

**DOI:** 10.1101/2025.07.04.663170

**Authors:** Lucas F. Jansen Klomp, Xinqi Yan, Rebecca R. Snabel, Gert Jan C. Veenstra, Hil G.E. Meijer, Janine N. Post

**Affiliations:** Department of Applied Mathematics, Faculty of Electrical Engineering, University of Twente, Drienerlolaan, Enschede, 7522NB, The Netherlands; Developmental BioEngineering, Faculty of Science and Technology, University of Twente, Drienerlolaan, Enschede, 7522NB, The Netherlands; Department of Molecular Developmental Biology, Radboud Institute for Molecular Life Sciences, Faculty of Science, Radboud University, Houtlaan, Nijmegen, 6525XZ, The Netherlands

**Author notes:** Contributing authors.

**Keywords:** Multi-omics, time series, ODE modelling, data-driven, mechanistic modelling, Gene Regulatory Network

## Abstract

**Background:** Understanding time-dependent intracellular processes, such as cell differentiation, is key to developing new therapies for a wide range of diseases. Models that connect transcription factor activity to dynamic expression patterns are rare, despite the increased availability of time-series data.

**Results:** To identify key regulators of time-dependent biological processes, we present the pipeline DANSE: **D**ynamics inference **A**lgorithm on **N**etworks **S**pecified by **E**nhancers. Starting from multi-omics data, our pipeline constructs a data-driven mechanistic transcription factor (TF) network and subsequently defines a dynamic model based on this TF network. The combination of a TF network and a mechanistic model allows for the identification of a small set of key transcription factors predicted to drive the modelled biological process. We showcase the result of our pipeline by applying DANSE to two different datasets that describe iPSC differentiation.

**Conclusions:** Models constructed using DANSE suggest testable hypotheses for the perturbation of gene expression, for example, knockdown or overexpression, that influence cell fate. In this way, DANSE is a powerful tool for generating novel hypotheses in a data-driven manner that take into account the dynamic nature of multi-omics time series data.

## 1 Background

Time-dependent intracellular processes, such as cell differentiation, are governed by complex interactions between signalling pathways and transcription factor (TF) networks. In recent years, time-series multi-omics data have provided a way to elucidate these temporal processes in the cell. Many methods of analysing these data have been proposed, including differential expression analysis, cell annotation and clustering, as well as forecasting new time points from existing data [1, 2]. However, their outputs do not directly provide mechanistic insight into which key genes control temporal changes in expression levels. To gain such mechanistic insights, dynamic models based on gene regulatory networks (GRNs) have been proposed [3–5]. The construction of such dynamic models first requires the inference of a suitable gene regulatory network. This task requires specialised inference methods that focus specifically on mechanistic connections between genes.

In literature, many approaches have been proposed to infer gene regulatory networks from multi-omics data [6–8]. Originally, these approaches focused on inferring networks based on bulk RNA-seq data [9, 10]. Recently, the focus has shifted to using multi-omics data [11–13] for GRN inference. Data collected at intracellular levels other than the mRNA level are often used to identify relationships between transcription factors and cis-regulatory elements, such as enhancer or promoter regions [6]. The benefit of these approaches is that there is more evidence for each inferred connection, leading to a clearer biological interpretation of the inferred networks. Common methods for inferring GRNs from multi-omics data include SCENIC+, which focuses on retrieving sets of a transcription factor (TF) and their targets from single-cell multi-omics data [14]. The method GraNIE infers both TF-peak and peak-gene links from RNA-seq and ATAC-seq data [12]. In this work, we consider the network inference method ANANSE [11] that uses a prediction of TF-enhancer binding to infer a GRN. The method ANANSE is particularly interesting for investigating time-dependent data, since the method is built to find key transcriptions involved in trans-differentiation. As a validated method to determine key genes distinguishing different cell fates, ANANSE is a promising tool for the construction of cell-type specific networks throughout cell differentiation.

Gene regulatory networks are often used as a basis for dynamic models, which provide additional mechanistic insight. In some inference methods, a gene regulatory network is inferred jointly with its dynamics [15–18]. However, these methods often focus on the inference of the gene regulatory network, rather than on dynamics. Other works describe how a predefined GRN is converted to an ordinary differential equations (ODE) model that mimics experimental data by fitting kinetic parameters [3, 4]. Some ODE models have been used to describe and identify intracellular mechanisms in cell fate decisions and signalling pathways [19, 20]. Though ODE modelling forms a natural paradigm for exploring gene expression over time, current methods are often focused on modelling known biological processes, reducing the ability to find key novel markers.

Here, we present DANSE: **D**ynamics inference **A**lgorithm on **N**etworks **S**pecified by **E**nhancers. In this pipeline, we use TF networks inferred using ANANSE to select key TFs from data and to infer a baseline gene regulatory network that indicates probable connections between the selected transcription factors. Subsequently, we infer a dynamic model based on this gene regulatory network, finding a minimal network that mimics the gene expression dynamics of modelled TFs. Through this process, we refine the inferred gene regulatory network and construct a mechanistic model of the temporal process described in the experimental data. This model provides direct mechanistic insights into possible drivers of the temporal process through varying parameters in the model. While many current popular analysis methods result in large sets of important TFs, the analysis of our dynamic models results in the identification of prioritised short list of TFs based on temporal dynamics, which are interesting for conducting knockdown or overexpression experiments in a wet-lab setting. We tested the performance of our pipeline with two multi-omics datasets: one describes pluripotent stem cells differentiating toward chondrocytes, while another dataset describes differentiation toward epicardioids [21–23].

## 2 Methods

The DANSE pipeline requires several inputs. First, multi-omics data at various time points are used to build the data-driven model. Second, the desired number of activating connections, inhibiting connections and transcription factors should be indicated for the generated model. Finally, a choice for the output TFs used to determine important transcription factors in the network is required. Based on these inputs, DANSE provides the functionality to build a computational model for the biological process described by the data, along with the capability to suggest novel drivers underlying these processes. Below, we describe and discuss all the choices made in this process of constructing a data-driven model based on multi-omics time series data. For a flowchart describing the workflow when using DANSE, we refer to Figure 1.

**Fig. 1:**
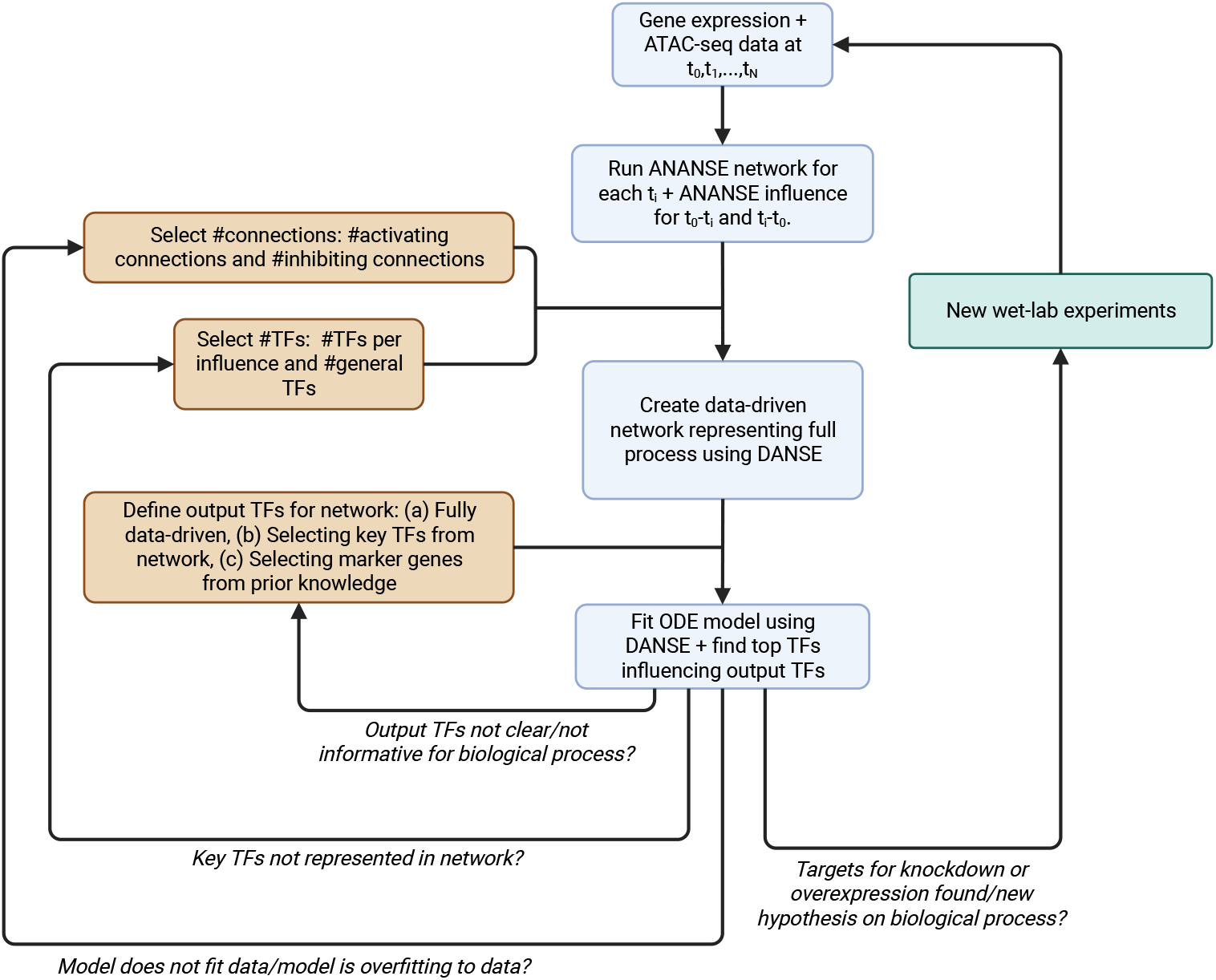
Flowchart for using our proposed pipeline, DANSE. The required input data is time series multi-omics data of an intracellular biological process. Three user-defined inputs are needed: the number of transcription factors and interactions desired for the computational model and the choice for output TFs used to rank TFs in the network. The DANSE pipeline then constructs a computational model based on the input. Subsequently, if the model needs adjustment, the choice for the number of TFs or the number of interactions can be adjusted. If not, the output of the model can be used to suggest knockdown or overexpression experiments, along with predicting key drivers underlying the considered process. If desired, new computational models can be generated based on the results of these experiments.

### 2.1 Network construction

The first step of our pipeline is the construction of a suitable gene regulatory network on which to base the dynamic model. Such a suitable network needs to include connections that are mechanistically realistic. Moreover, such a network should consist of somewhere between 10-50 nodes, as networks with too many nodes may be hard to interpret, while too few nodes may lead to missing essential regulators. Finally, the constructed network should include both activating and inhibiting connections. Including inhibiting connections in the network allows for the construction of more types of feedback loops in the network, which are key to describing specific behaviour using dynamic models (for example, bistability or oscillations).

To construct such a network, we first construct fully connected networks for each available individual time point using ANANSE, using RNA-seq data and ATAC-seq data collected at those time points as input (Figure 2b). We then use the influence score implemented in ANANSE to identify TFs that change significantly in importance between time points [11]. This influence score is based on changes between time points in network connectivity and expression levels. Hence, it is a measure that identifies TFs that are important in one time point compared to another. By including TFs with a high influence score in our model, we aim to identify TFs that have an important role in one or more specific stages during differentiation.

**Fig. 2:**
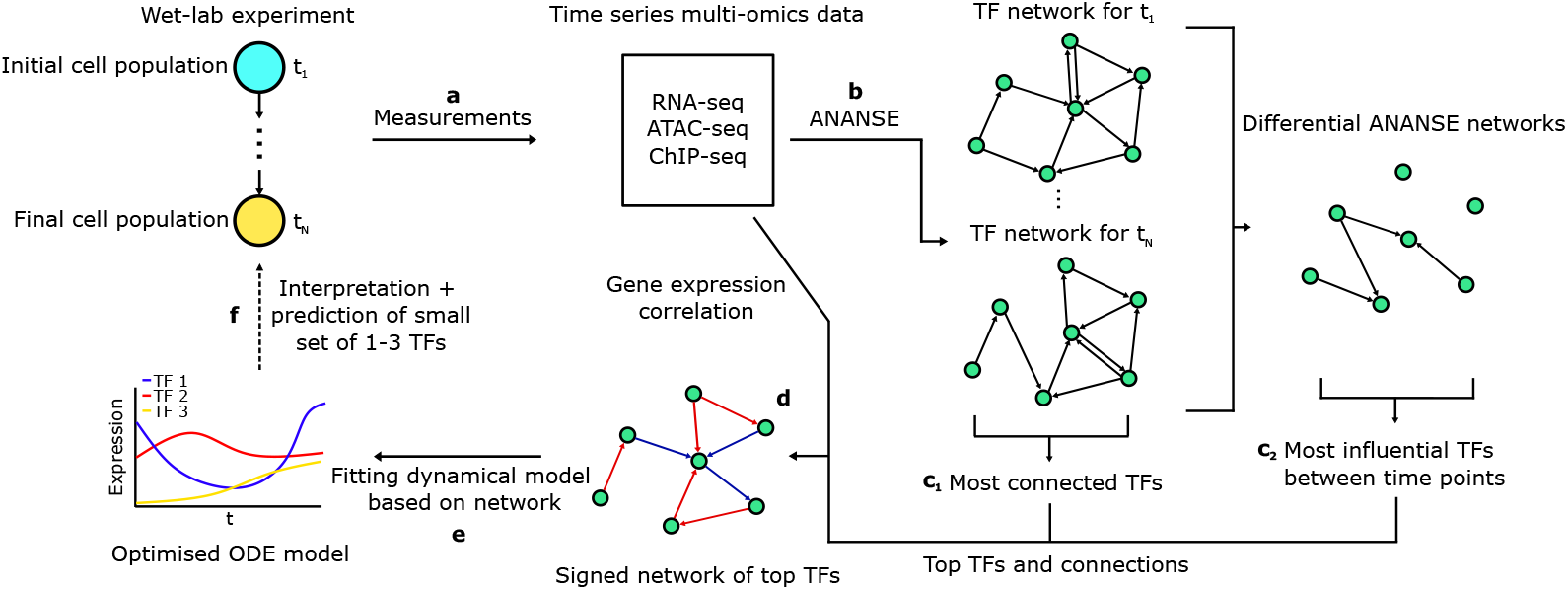
Overview of the DANSE pipeline for making dynamic models based on timeseries multi-omics data. multi-omics data in the form of gene expression data, ATAC-seq data and optionally ChIP-seq data is used to infer a GRN that is suitable for dynamic modelling using the GRN inference tool ANANSE. Subsequently, an ordinary differential equations model is defined based on this network and parameters in this model are optimised, leading to a minimal model that describes the dynamics found in the original experiment (a schematic simulation of TF expression is shown here). Insights and mechanisms found in this computational model can be used to inform decisions on new experiments, leading to a modelling cycle.

To select TFs for our network, we compare the network of each individual time point with the network for the first time point in the differentiation experiment. In this way, we aim to identify TFs that best explain the differences between these cell states. We computed the ANANSE influence score both forward and backward in time. In this way, we identified TFs that are more important at each later time point compared to the first time point and TFs that are more important at the first time point compared to each later time point. For each comparison, we include the top genes ranked by influence score in our network (Figure 2c_2_, Figure B3). This yields a set of genes per time point after the first time point. In order to include TFs that are important throughout the entire differentiation process, we also included in our network topology the top TFs ranked by the average number of network connections over all time points (Figure 2c_1_). Finally, we remove all duplicate genes from our set of selected TFs.

Connections between the TFs are selected based on the static networks constructed for each of the considered time points (Figure 2d). To accommodate potential context-dependent effects based on the composition of regulatory elements, we designed our model such that TFs could have variable effects on target genes. Therefore, we first sign each possible connection, defining them as activating or inhibiting based on the correlation between gene expression of the source and target TFs. We then select the top activating and inhibiting connections as defined by the maximum of the interaction scores over all time points for which a network is made. The interaction score *A*_*ij*_ between TFs *i* and *j* is the connection strength defined using ANANSE:

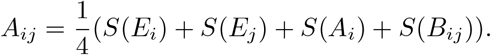

Here, *E*_*i*_ denotes the expression of TF *i, A*_*i*_ denotes genome-wide activity of TF *i* and *B*_*ij*_ denotes inferred binding probability of TF *i* to enhancer regions close to the gene associated with target TF *j* [11]. The function *S* scales each value to the interval [0, 1]. We use a different interaction score for the inhibiting connections (as defined by a negative correlation in gene expression), where for the expression of the target TF *j*, a low expression level is preferred, so:

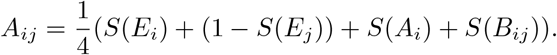

We note that binding of TFs to enhancer regions is generally associated with upregulation of target genes. However, silencers, the inhibition-associated counterparts to enhancers, are not as well documented. Therefore, we have chosen the above formulation to be able to identify potential inhibiting connections which are key for modelling, but we emphasise that these connection strengths are not mechanistically accurate.

Based on these interaction scores, we select the top *n*_*a*_ activating connections and the top *n*_*i*_ inhibiting connections. Subsequently, we only keep those nodes in the network that have at least one incoming activating connection, removing all others, and repeat until no such nodes remain in the network. Hence, in our pipeline, the potential nodes in the network are selected first, after which the connections are iteratively added to the network. Because we remove nodes that get no activating input after selecting connections, the number of nodes in the final TF network depends on the number of activating connections chosen. The selection of the number of connections should be based on the resulting dynamical model. We comment on this in more detail when presenting the results of the application of DANSE.

### 2.2 Dynamic modelling

Based on the constructed TF network, the aim of the DANSE pipeline is to describe time-series data and explore mechanisms underlying the data. To do so, we formulate an ordinary differential equations model that describes the change in expression of each transcription factor *x*_*i*_ (Figure 2e). Each TF decays naturally over time and its activation or inhibition by other TFs is modelled by a Hill function. The ODE system is given by:

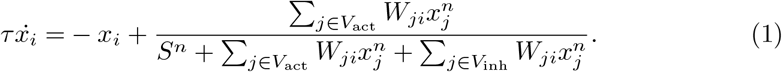

In this system, the weight matrix *W*_*ji*_ denotes how strongly TF *j* influences *i*. As such, the matrix *W* describes the connection strength for the entire network. If there is no connection between TF *i* and *j, W*_*ij*_ = *W*_*ji*_ = 0. The sets *V*_act_ and *V*_inh_ denote the sets of TFs that activate and inhibit TF *i* respectively. The parameter *S* denotes the half occupation value, and *n* is the Hill coefficient. We set *S* = 1 and *n* = 3, leading to a reasonably steep sigmoid shape. Such a steep sigmoid leads to easier representation of bistability within the system. The parameter *τ* describes the time scale of the full system.

Models based on Hill functions are widely used in literature to model gene regulatory networks [24, 25]. The specific form of the Hill function used in our pipeline is also described in literature [19] and has several desirable properties. First, because the Hill function is bounded between 0 and 1, the activity of a modelled TF will stay between 0 and 1 if its initial conditions are between 0 and 1. This allows for modelling normalised and scaled RNA-seq data. Second, this formulation allows for large effects of a single activating or inhibiting input in the absence of other influences. Because of this, our formulation is less sensitive to differences in the amount of incoming connections for each node. We use a single connection strength for each connection in the network, and do not adjust the connection strength according to changing chromatin accessibility over the full temporal process.

### 2.3 Parameter optimisation

We optimise the weight matrix *W* in the ODE model to fit time-series expression data of the modelled TFs (Figure 2e). Given data *D*_*i*_(*t*) denoting the expression of TF *i* at time point *t* and a solution *ϕ*(*t, W*) ∈ ℝ^*N*^ for a given *W* to the system of *N* ODEs given by Equation 1 with *ϕ*_*i*_(0, *W*) = *D*_*i*_(0), we optimise *W* by numerically minimising a loss function given by:

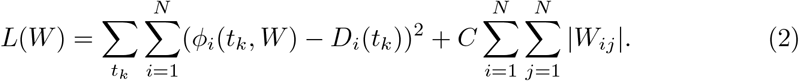

Here, *ϕ*_*i*_(*t, W*) denotes the *i*th element of *ϕ*(*t, W*). The first term in this loss function denotes the squared error between the data and the simulation. For each connection between TFs *i* and *j* that is not part of the full inferred gene regulatory network, we fix *W*_*ij*_ = 0; these elements of the weight matrix are not optimised. We note that *W* is static throughout the simulation of the model.

The second term in the loss function is an L1 regularisation term. This term penalises any nonzero elements in the weight matrix. The penalty is multiplied by a scalar *C* ∈ ℝ. By setting a low value for *C, C* = 0.005, we ensure that connections in the gene regulatory network that do not contribute significantly to fitting experimental data are removed. On the other hand, taking a low value for *C* ensures that a good fit of the data is prioritised. In turn, the L1 regularisation term will only remove connections that do not add anything to the fit of the model, or connections that have the same function as other connections in the network. In this way, our optimisation procedure looks for a minimal network structure that simulates the experimental data with an added restriction that any connection included in the model must be feasible based on the given experimental data, based on the original constructed TF network.

We optimise the loss function using the gradient-based optimiser ADAM with learning rate 0.1 [26]. We note that this is a high learning rate, but we have found that lower learning rates do not yield better fitting results or more stable training behaviour in our test cases.

A choice made during the preprocessing of expression level data is how this data should be scaled to give as input for the parameter fitting procedure. DANSE takes normalised expression levels as input, and scales the data so that expression levels for all genes fall within the interval [0, 1]. That is, we compute the scaled expression level of gene *i*, denoted by *X*_*i,t*_, from the normalised expression level 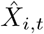 by:

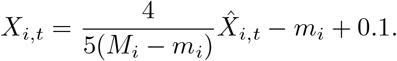

Here, *M*_*i*_ and *m*_*i*_ are the maximum and minimum expression levels for gene *i*, respectively. This scaling procedure scales the expression of all genes between 0.1 and 0.9, where the maximum expression level of a gene is assigned a scaled expression of 0.9, and the minimum expression level is assigned a scaled expression of 0.1. From a modelling perspective, this ensures that there is ample variation in gene expression for each modelled gene, and therefore the model cannot “ignore” particular genes in the network. As only highly active TFs and TFs with high changes in activity are included in the model, stable lowly expressed genes are not highlighted in an undesirable way using this procedure. From a biological perspective, genes with low expression levels may still have a significant influence on the rest of the system, hence why alternative methods such as scaling over all genes are not viable options.

A choice made prior to running the DANSE pipeline, the choice of the number of connections in the model, can be based on the results from fitting the model. Specifically, the aim is to select the least number of connections necessary to obtain a reasonable fit of the data, hence aiming to avoid overfitting of the data (here defined as parsimony) or having too many options for recreating the data using the selected network. A selection of too many connections may in some cases lead to biologically unfeasible behaviour such as oscillations. To determine this threshold, we recommend to optimise the model multiple times for different choices of the number of activating connections *n*_*a*_ and inhibiting connections *n*_*i*_. In this way, a grid search is performed for the choice of the number of connections in the network. We consider the average value of the loss function for all nodes in the network to be an indicator of the fitting performance:

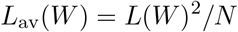

where *N* denotes the number of nodes in the network for that particular optimization. If this metric drops significantly when adding new connections to the model, this is an indication that the fit has performed well. We describe other criteria for selecting these parameters when discussing the results for our case studies.

### 2.4 Identification of key transcription factors

As a method of obtaining novel insights based on the constructed dynamic model, we focus on the discovery of key transcription factors (Figure 2e). The metric we use to determine the importance of a TF relies on how much upregulation or downregulation of this TF influences the expression of selected key TFs: transcription factors associated with an on-target cell fate. Upregulation and downregulation of TFs is modelled by modifying the right-hand side of the ODE model for a TF *i* that is upregulated or knocked down:

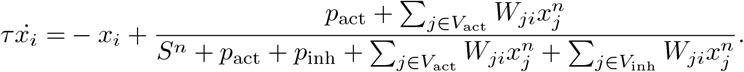

Here, *p*_act_ and *p*_inh_ are the controlled variables used to indicate how much TF *i* is upregulated or knocked down. When *p*_act_ = *p*_inh_ = 0, there is no change compared to the original model. When *p*_act_ *>* 0, *p*_inh_ = 0, transcription factor *i* is upregulated. On the other hand, when *p*_inh_ *>* 0, *p*_act_ = 0, transcription factor *i* is downregulated. In this controlled case we do not consider the case where *p*_act_ and *p*_inh_ are both non-zero. Specifically, we modify one parameter *p* so that

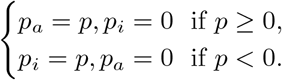

We then define a set of transcription factors *T* = {*i*_1_, *i*_2_, …, *i*_*n*_} of transcription factors *i*_*k*_ that are known or inferred to be markers for the modelled biological process associated with the desired cell fate, selected from the TFs present in the inferred TF network, from data or from prior knowledge. For each other TF in the network, we then find:

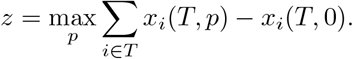

This maximum *z* is a metric for how much the regulated transcription factor influences the set of transcription factors *T*. In this metric, we focus only on upregulation of the target genes in *T*, since the selected set of known transcription factors are assumed to be desirable for a specific cell fate. By calculating our metric for each node in the network, we can obtain a list of the main transcription factors that influence specific markers according to the computational model constructed using DANSE. Our method will present a prioritised set of TFs of interest, eliminating the need to sift through large sets of identified transcription factors. We highlight here that the novelty of our selection method is that it takes the temporal dynamics into account, which differs from methods using only one time point. Moreover, since we have the underlying computational model, we can subsequently identify an underlying mechanism explaining how this particular TF may upregulate the set of selected target transcription factors. In some cases, these effects can be indirect and are only clearly visible through the introduced dynamics. In this way, the DANSE pipeline provides a novel way of generating hypotheses based on multi-omics time series data. We highlight here that the difference between classical gene expression analysis and our method is that our method of finding key TFs is explicitly based on the constructed mechanistic dynamic model.

We provide three options to choose the set of key transcription factors *T*. First, TFs can be selected in a fully data-driven way. Here, the TFs that return most often in influence scores calculated using ANANSE going forward in time are added to the network as output nodes. If these nodes are not already in the TF network, the top two incoming activating connections and the top two incoming inhibiting connections are added based on the ANANSE interaction score. Second, TFs can be selected directly from the final TF network by hand. This is a good option if TFs are included in the network which have a well-known function within the modelled biological process. Finally, TFs can be selected from prior knowledge. Similar to the data-driven option, the top two incoming activating and the top two incoming inhibiting connections to these TFs are included in the network. This option is most relevant if you want to investigate the regulation of particular marker genes known from other work.

Since our model optimisation procedure yields slightly different optimised models in multiple runs, we suggest running the optimisation multiple times and recording the importance measure *z* for each TF in the network for each individual fit. By measuring statistics of the importance of the TFs on target genes of interest over different runs, it is possible to quantify the robustness of our method. We showcase this procedure in the two case studies in the Results section.

## 3 Results

We illustrate the use of the DANSE pipeline by applying it to two publicly available datasets. Specifically, we ask whether it is possible to obtain novel biological insights into pluripotent stem cell differentiation using our pipeline. We consider a dataset describing chondrogenesis and a dataset describing epicardioid differentiation.

### 3.1 Chondrogenic differentiation

To explore chondrogenesis, we have applied DANSE to a dataset generated from a chondrogenesis experiment conducted by Kawata et al. [21]. From this experiment, microarray and bulk ATAC-seq data is available for days 0,2,3,4,5 and 9 during differentiation. The differentiation relied on a combination of two compounds, a WNT activator and retinoic acid, to differentiate iPSCs towards chondrocytes. The raw data has been preprocessed before being used as input for DANSE. For details on this preprocessing procedure we refer to Section A.

To construct a model using DANSE for this dataset, we selected the top 5 genes per comparison between day zero and a later time point, and selected the top 12 generally important transcription factors. We chose to include the top 110 activating connections and the top 30 inhibiting connections in the network. This was based on rerunning the pipeline for various choices for the number of activating and inhibiting connections (Figure B1). Adding fewer connections led to a good fit, but then few nodes are included in the network. At the threshold of adding 110 connections, the fit of the model improved noticeably compared to adding fewer activating connections.

By running the DANSE pipeline, we obtained a dynamical model for the chondrogenesis process that mimics the gene expression profiles in the data (Figure 3B). The constructed data-driven TF network contained various important known TFs (Figure 3A). The transcription factors RXRA, RARG and TCF3 are directly related to retinoic acid signalling and WNT signalling, the two stimuli applied in the protocol described by [21]. In addition, SOX8 is an important gene in chondrocytes and is related to the chondrocyte master transcription factor SOX9 [27], and GLI3 is associated with hedgehog signalling and has a known role in chondrogenesis [28, 29]. Hence, the model also included some genes that have less well-defined roles within chondrogenesis, such as SREBF1 and BCL11A.

**Fig. 3:**
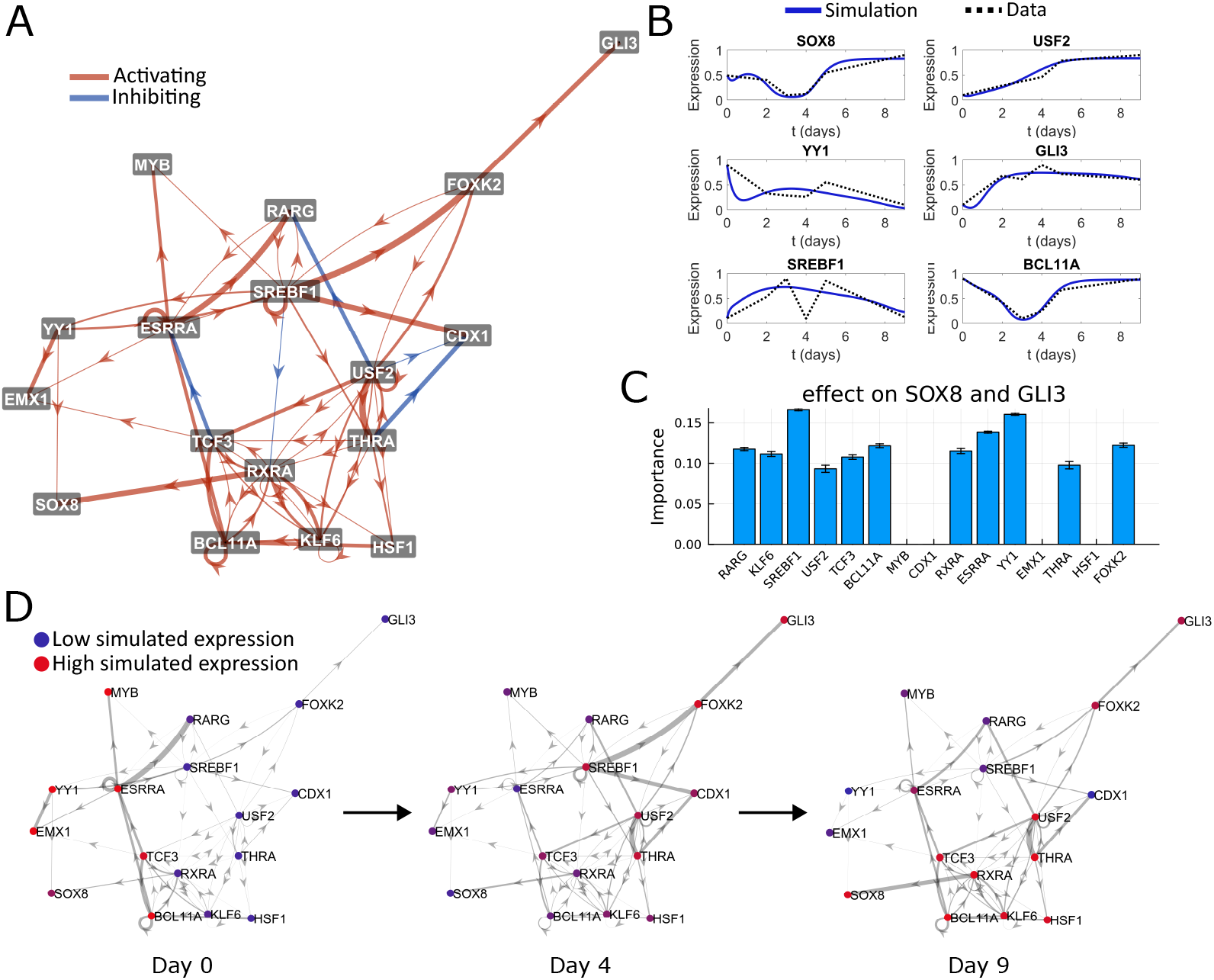
Computational model constructed using DANSE based on the data from Kawata et al. [21] **A**. Constructed TF network based on the multi-omics data describing the full differentiation process. Here, activating connections are indicated in red and inhibiting connections are indicated in blue. The thickness of each connection indicates the strength of that connection in the fitted dynamic model. The position of the nodes is not related to the model **B**. Simulations of the fitted dynamic model, showing a qualitative match between model simulations and gene expression level data. **C**. Calculated importance measures for all TFs showing which TFs in the network have the most effect on SOX8 and GLI1 expression when knocked down or overexpressed. **D**. Visualisation of expression levels over time in the TF network. The colors of the nodes indicate simulated expression levels of the TFs at the depicted time points. The thickness of the edges depicts how much signal is going through each connection, defined as the activity of the source node multiplied by the fitted connection strength.

To obtain potential targets influencing the chondrogenesis process, we manually identified SOX8 and GLI3 as factors present in the TF network associated with on-target chondrocyte differentiation. We then asked which other factors in the constructed model are promising targets for modulating the expression of these selected TFs. Our data-driven model suggested that many TFs in the network have some effect on the expression of SOX8 and GLI3. However, SREBF1, YY1 and ESRRA are the most promising targets following our importance metric (Figure 3C). Two of these genes have a known function in cartilage differentiation: the gene YY1 is known to play a role in mesenchymal stem cell differentiation towards cartilage [30], while ESRRA has been shown to directly regulate SOX9 in zebrafish. Moreover, knockdown of ESRRA has been shown to induce expression of osteoarthritic markers in human chondrocytes [31, 32]. Less is known about the function of SREBF1 in chondrogenesis, making this a potential new target for knockdown or overexpression experiments. Our result persists locally if small changes are made in the number of selected activating or inhibiting connections (Figure B4). If larger changes are made to the network structure, other genes can take the top spot in terms of the importance measure, but SREBF1 is highly ranked for alternative choices of the number of connections in the network.

Novel biological hypotheses can be formulated based on the identification of these important transcription factors in our computational model. Taking SREBF1 as an example, the fitted TF network suggested that GLI3 is affected indirectly by SREBF1 through the activation of FOXK2. Additionally, SREBF1 has an effect on the regulatory network steering SOX8 expression through activating THRA and inhibiting RXRA. As such, the application of DANSE resulted in testable hypotheses on the workings of intracellular dynamics, which can be validated in wet-lab experiments through knock-down and overexpression experiments. While the TFs included in the computational model were based on the influence score computed using ANANSE, the order in which TFs are ranked using DANSE is different from when ANANSE is used directly. This shows how the construction of the computational model lead to novel insights into the gene regulatory network structure. In general, the focus of many existing methods is to find genes that are differentially expressed or have different activity across cell types, whereas our method aims to find the regulators of these differentially expressed genes or differentially active transcription factors.

### 3.2 Epicardioid differentiation

To illustrate the use of our pipeline for exploring single-cell data, we applied DANSE to a single-cell multi-omics dataset which describes iPSC differentiation to epicardioid cells [23]. The data describes the formation of self-organizing human pluripotent stem cell-derived epicardioids that display patterning of the epicardium and myocardium. scRNA-seq and scATAC-seq data were collected on day 2,3,4,5,7,10 and 15 during the differentiation experiment. Details on data preprocessing for use in DANSE can be found in Section A.

To construct the dynamic model describing the epicardioid differentiation process, we first selected the number of activating and inhibiting connections for the model in order to obtain a good fit of the model to the data. From the optimization results in (Figure B2), the model performed well when 140 activating connections and 30 inhibiting connections were included. While the model also performs well for some lower values of the number of activating and inhibiting connections, these choices result in very few genes being included in the model. We then zoomed in to explore other selections for the number of activating and inhibiting connections in the neighbourhood of this choice and found that there is a cut-off in the value of the loss function near 140 activating connections (Figure B2). Therefore, we have picked 140 activating and 30 inhibiting connections to be used for our model.

In the constructed TF network we identified various known relevant transcription factors in cardiac development (Figure 4A). For example, TBX3 is known to repress TBX5, which is essential for early heart tube development and activates the chamber formation [34]. MEIS2 is essential for cardiac neural crest development [35]. Homeobox genes (HOXB2/4) contribute to patterning of cardiac progenitor cells and formation of the great arteries [36]. PBX1 functions in distinct regulatory networks to pattern the great arteries and cardiac outflow tract [37]. Simulations of the model constructed using DANSE mimic the experimental data (Figure 4B).

**Fig. 4:**
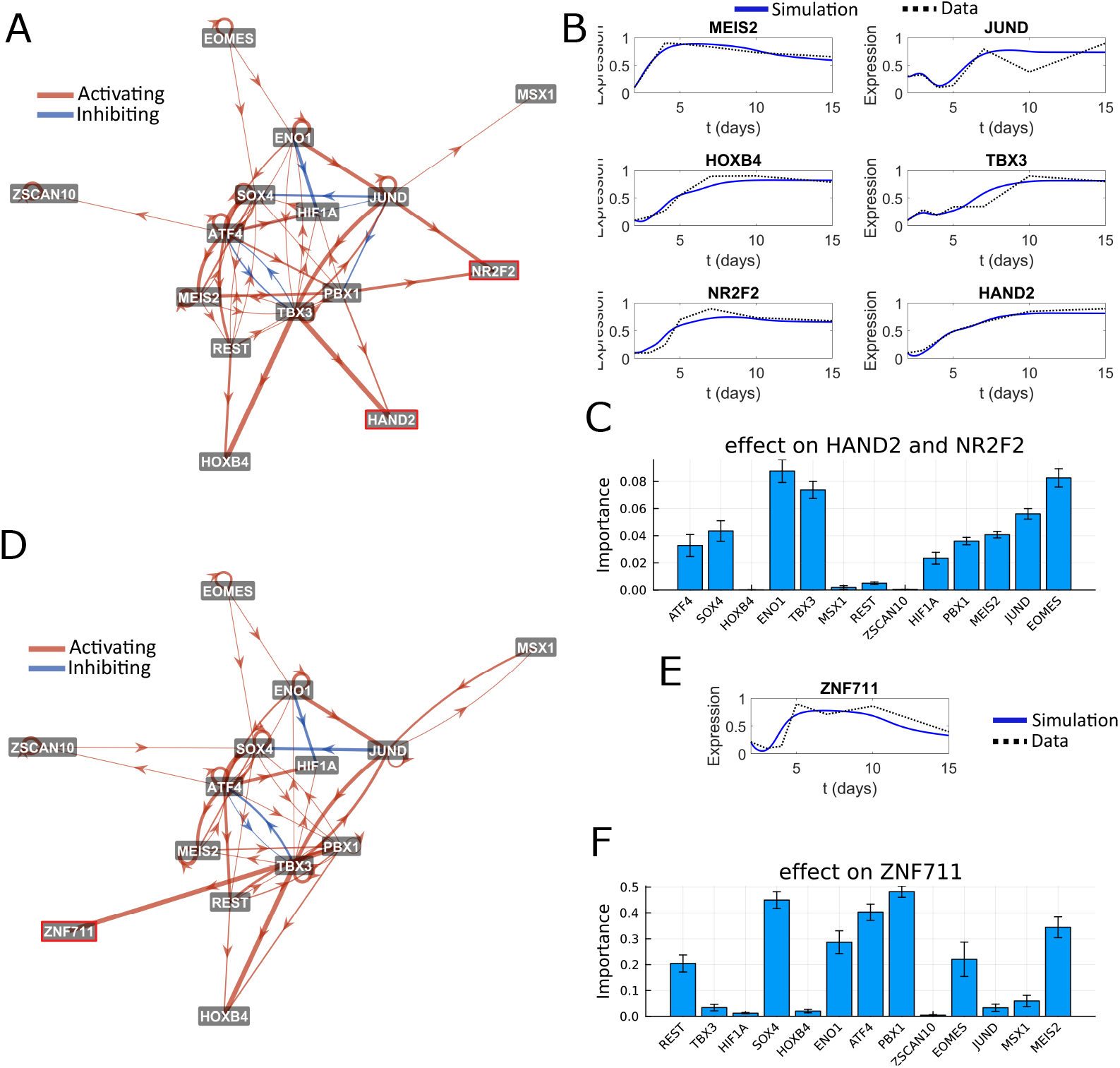
Dynamic model of epicardioid differentiation based on data from Meier et al. [23] **A**. Network topology of the model with two additional nodes: HAND2 and NR2F2, selected using our data-driven approach for finding marker genes. **B**. Simulation results of important TFs in the model including HAND2 and NR2F2. Blue lines denote simulations, while dashed lines denote experimental data. We see that the simulation fits the experimental data well. **C**. Our model identifies ENO1, EOMES and TBX3 to be the most important TFs to regulate HAND2 and NR2F2 expression. **D**. Network topology of the model with one additional node: ZNF711, selected from prior knowledge based on [33]. Simulation of ZNF711 expression fits the experimental data well. **F**. Our model identifies PBX1, SOX4 and ATF4 to be the most important TFs to regulate ZNF711 expression.

To identify key TFs for genes of interest, we showcase the data-driven option and the prior knowledge option of the DANSE pipeline for selecting marker TFs, after which novel potential regulators can be found. When applying the data-driven option, HAND2 and NR2F2 were selected as marker TFs based on the ANANSE influence score. This option is purely data-driven and not biased based on prior knowledge. From a biological perspective, these markers selected based on the data have clear roles within epicardioid differentiation. HAND2 is a well-known marker for heart tube cells and right ventricle cardiomyocytes and its function in second heart field progenitors is essential for cardiogenesis [38]. The other selected marker gene, NR2F2, also known as COUP-TFII is an atrial cardiomyocyte marker [39]. As neither HAND2 nor NR2F2 was included according to the top connections in the original model, we added them during the construction of the network. To do so, we add the two most likely activating and two most likely inhibiting connections to the target genes to our TF network. These most likely connections are identified by the highest interaction score over all measured time points. We then asked which genes are most likely to affect NR2F2 and HAND2 activity.

Our model identified ENO1, TBX3, and EOMES as potential regulators of HAND2 and NR2F2 during epicardioid differentiation. The transcription factor EOMES is known to be important during early iPSC differentiation, and it has been shown that EOMES induction can induce cardiac differentiation [40]. On the other hand, TBX3 has been shown to induce differentiation towards pacemaker-like cells, a cell type distinct from the targeted epicardiac cells [41, 42], but has also been shown to cooperate with HAND2 directly [43]. The transcription factor ENO1 has a known role in cardiac contraction and has recently been identified to have a protective role against heart failure [44, 45]. However, its effect on iPSC differentiation towards epicardioid cells is not clear from current literature. These results show the capability of DANSE to recover both known drivers of cardiac differentiation, as well as potential targets for which the role in cardiac differentiation is not as well-described in current literature.

When looking at the most important TF calculated from different choices for the amount of activating and inhibiting connections, we found that the three identified TFs are also inferred to be important regulators of HAND2 and NR2F2 for other choices of the number of activating and inhibiting connections. Even though the most important TF does change depending on the number of connections in the model, ENO1 is recovered frequently as a TF that potentially regulates HAND2 and NR2F2 (Figure B5). This result shows some robustness of our method to recover the most important TF from the model.

In addition to the data-driven approach, we illustrate how to use our pipeline when the selection of marker genes is purely driven by prior knowledge. In particular, ZNF711 has recently been identified as a safeguard for cardiomyocyte commitment [33], and it is therefore interesting to ask which TFs may regulate ZNF711 expression. To investigate this, we added ZNF711 to the data-driven TF network following the same procedure as when adding HAND2 and NR2F2. We then asked which TFs in the data-driven network are most likely to influence ZNF711 expression. We identified PBX1, SOX4 and ATF4 as potential regulators of ZNF711. These three TFs are also recovered as potential regulators of ZNF711 for various alternative choices of the number of connections in the network (Figure B6). Especially interesting is the role of PBX1, as this TF is predicted to directly regulate ZNF711 in our computational model. While we could not find evidence of this direct regulation in literature, PBX1 has previously been found to collaborate with HAND2 in limb development [46] and has been found to directly regulate NR2F2 during steroidogenesis [47]. The combination of our model results and the known function of PBX1 in cardiac differentiation makes it a promising target for optimising differentiation experiments toward cardiomyocyte cells.

## 4 Discussion

In this work we present DANSE, a pipeline that can be used to construct a dynamic model based on time-series multi-omics data without explicitly providing prior knowledge-based input such as an initial gene regulatory network. We have illustrated the use of this pipeline by applying it to bulk data collected during a chondrogenesis experiment and single cell data collected during a cardiogenesis experiment [21, 23]. The constructed TF networks contain transcription factors known to be important for both processes, and the resulting dynamic models mimic experimental gene expression data. Subsequently, we have shown that our pipeline leads to the identification of interesting targets for upregulation and downregulation. Our method of defining important transcription factors looks beyond the local role of individual nodes, instead focusing on the role of transcription factors within the full TF network. Although ATAC-seq data is always required, we show through the two applications of our pipeline that DANSE is applicable to both RNA-seq data and microarray data.

Beyond confirming known regulators, DANSE serves as a hypothesis-generating tool for biological discovery. It enables researchers to ask targeted, mechanistic questions that are central to understanding cell fate decisions. For example, based on the constructed model, it is possible to investigate which transcription factors drive a cell fate transition over time or what upstream regulators must be active to induce these key drivers. The pipeline also supports in silico perturbation experiments, allowing biologists to explore how changes in TF expression affect the network as a whole, opening the door to using experimental validation using tools like CRISPR-Cas. Through the executable nature of the constructed computational models, our pipeline goes beyond what is possible with conventional methods such as pathway analysis [48, 49]. By translating multi-omics data into interpretable, dynamic models, DANSE creates a bridge between complex data and testable biological hypotheses and ultimately, a path toward predictive and programmable biology.

In recent years, the advent of single-cell sequencing has led to a drastic increase in detail of generated data. Pseudo-time trajectory inference is a method commonly used to describe temporal processes based on single-cell sequencing data [50, 51]. Through the resulting pseudo-time trajectory, branching points and other interesting features of a dataset can be explored. These methods work well to describe the data, but do not directly provide mechanistic insights. While our pipeline is built for and initially tested on bulk data, it is also applicable to single-cell data, as shown in the application to the epicardioid differentiation experiment [23]. After an appropriate clustering of the single-cell data, TF networks can be constructed for each individual cluster using ANANSE [13]. Subsequently, a procedure similar to the one described in this work could be used to select key TFs from each cluster and build a full TF network. The ODE model posed on this network can be fitted to pseudo-time trajectories, resulting in a mechanistic model describing single-cell data.

To run the DANSE pipeline, an initial choice of the number of activating and inhibiting interactions as well as the desired number of TFs for the network is required. In a practical use case, this means that the pipeline should be run multiple times to account for the effects of these hyperparameter choices. For the choice of TFs, ANANSE influence plots can be investigated to see which TFs are included in the network topology. Based on these plots, a choice for the cut-off value for the number of TFs can be made. For the choice of the number of activating and inhibiting connections, we suggest running DANSE multiple times for different choices of the number of activating and inhibiting connections. More connections should be added to the model if the resulting models do not match trends in the experimental gene expression data. Fewer connections should be selected if the model shows biologically unfeasible behaviour (for example, oscillations in a known non-oscillating system) or very different mechanisms for each run fit of the model are found. Moreover, the final value of the loss function for the different fitted models serves as a metric for the performance of the model; if this value drops significantly when adding connections, it is likely that adding these connections is a good decision. We note that it is not possible to automate these hyperparameter choices, since different input data lead to models with different characteristics. Therefore, we emphasize that these choices are a necessary modelling choice, while DANSE provides a tool to convert these modelling choices to a dynamic model along with a first analysis of the resulting model. In this way, DANSE should be seen as part of a modelling cycle, where the model is iteratively updated until a prediction is made, leading to new data which can be incorporated in the model. We refer to Figure 1 for an overview of the workflow using DANSE.

When selecting the hyperparameters necessary to run the DANSE pipeline, a concern is whether the model is overfitting to the data. We have found that while the top identified key TF does not change with every added or removed connection, the output of our pipeline does vary when considering wider ranges of hyperparameter choices. This reinforces the need for the setup of a modelling cycle, where the alternative hypotheses posed by different computational models can be validated in a wet-lab setting. We note that conventional methods of checking overfitting are not readily applicable to the models generated using DANSE, as only the small TF network and gene expression as sparse time points for these TFs is used to fit the model. Therefore, leaving out one data point as a “test set” is possible, but will result in removing so much information from both the network construction and time series data used for fitting the model that it becomes difficult to compare results. We have therefore chosen to focus only on model choices and hyperparameters to evaluate the performance of models constructed using DANSE.

In the current approach the interaction score is allocated a sign that reflects the predicted effect of a TF on the collective enhancer regions connected to a specific target gene. This interaction is activating or inhibiting, which means this model allows a TF to be both a repressor or activator in the same network, depending on which enhancers it is binding. From a mechanistic point of view, this is a simplification. With some exceptions, for example nuclear hormone receptors that act as an activator or repressor depending on the presence of the nuclear hormone ligand [52], transcription factors tend to be either activators or repressors and do not have both functionalities. However, the consequences of their perturbation can be highly complex, due to both indirect effects of the regulatory circuitry [53], and enhancer grammar and syntax [54]. Integrating complex regulatory input at the enhancer level can accommodate highly complex temporal or spatial expression patterns without invoking a variable activity of TFs [55]. On the other hand, enhancer grammar may involve a different number, orientation and spacing of motifs that can be bound by TFs, causing individual TFs to produce a context-dependent effect on a regulatory region. In DANSE, our approach is to model the TF’s effect on the regulatory regions in the context of that specific target gene - of which the expression is also taken along to calculate the interaction score (*E*_*j*_) - as a simplified model of the context-dependent effects of TFs on target genes.

A limitation of the DANSE pipeline is that we assume that connections as well as optimised connection strengths between TFs are static throughout a modelled process. This may not be the case in reality, where responses of cells to specific stimuli vary a lot in different cell types. A possible adjustment to the pipeline could be made, whereby a network is inferred for each individual time point. Computational models based on these individual networks could be linked together to form a multi-scale model of alonger temporal process.

Looking at the interpretation of the models built using DANSE, mechanistic modelling of (semi-)bulk data has restrictions resulting from the lack of granularity per measured time point. Therefore, mechanistic models built using our pipeline describe changes in gene expression of one population of cells over different time points, but cannot track multiple cell types, since the model is fit to one time series. This is a limiting factor in the case where single cell data is considered in which cells split into two trajectories. With our current implementation of the DANSE pipeline, it is only possible to construct a model for each of the trajectories individually. However, a model describing all possible trajectories may be desired. To achieve the identification of such a model, a potential approach is to redefine the procedure used for fitting the model to the data by simulating a population of cells, and using methods from optimal transport to compute the goodness of fit [56]. This probabilistic view of the fitting procedure may also help to overcome the fact that our current method does not take noise into account.

## 5 Conclusion

DANSE allows to go beyond static descriptions of gene regulation, and offers a framework for building dynamic, interpretable, models directly from multi-omics data. By integrating network inference based on transcription factor binding sites and transcriptomics with dynamic modelling, DANSE allows to go from observational data to mechanistic hypotheses in a data-driven manner. The ability to identify a minimal network of key regulators of cellular processes makes it a novel and powerful tool for uncovering both known and novel drivers of cell fate decisions. DANSE has the potential to transform the way we analyse time-series data in tissue development, regeneration and disease progression in a data-driven and objective manner.

## Declarations

### Ethics approval and consent to participate

Not applicable.

### Consent for publication

Not applicable.

### Availability of data and materials

All code for the DANSE pipeline is available on https://github.com/LJansenKlomp/DANSE.

The datasets analysed during this study are available in the NCBI Gene Expression Omnibus repository. The data for the chondrogenesis experiment described in [21] can be accessed using accession numbers GEO: GSE96036, GSE109172 and GSE132532. The data for the epicardioid differentiation experiment described in [23] can be accessed using accession number GEO: GSE196516.

We have used BioRender to create Figure 1. The license for this figure can be found at https://BioRender.com/44wye8n.

## Authors’ contributions

LJK was involved in the conception of the work, the design of the work, the analysis, the interpretation of data, the creation of software and drafting and revising the work.

XY was involved in the design of the work, the analysis, the interpretation of data, the creation of new software and the drafting and revising of the work.

RRS was involved in the design of the work and the revision of the work. GJCV was involved in the design of the work and the revision of the work.

HGEM was involved in the conception of the work, the design of the work, the analysis of the work and the revision of the work.

JNP was involved in the conception of the work, the design of the work, the analysis of the work and the revision of the work.

## Acknowledgments

We thank Ingrid Meulenbelt and Rick Mulders for the fruitful discussions as part of the SCI-MAP Bioinformatics meeting.

## Appendix A Supplementary Methods

In this section, we describe in detail the preprocessing steps used to get to the input data used for DANSE for the two computational models for chondrogenesis and cardiogenesis described in the main text.

### A.1 Preprocessing for chondrogenesis data

In the chondrogenesis experiment conducted by Kawata et al. [21], microarray and bulk ATAC-seq data was collected for early time points during differentiation from hiPSCs towards chondrocytes. Data was obtained from the NCBI Gene Expression Omnibus public repository (https://www.ncbi.nlm.nih.gov/geo/, accession numbers GEO: GSE96036, GSE109172 and GSE132532). The raw microarray data was log2 transformed, and subsequently normalised using limma (normalizeBetweenArrays) [57], accounting for the fact that the microarray chip was manufactured by Agilent. For each time point, three samples were available and we average them. These averaged expression levels were used as input for our network construction and parameter optimisation.

The deposited ATAC-seq data contains peak locations derived from the raw data. For each time point, we converted the data to a count matrix, a format supported for network construction using ANANSE [11]. The resulting count matrices are used as input for network construction using ANANSE.

### A.2 Preprocessing for epicardioid differentiation data

In the differentiation experiment towards epicardioid cells conducted by [23], single-cell data was collected during a differentiation experiment from hiPSCs towards epicardioid cells. Data was obtained from the NCBI Gene Expression Omnibus public repository (https://www.ncbi.nlm.nih.gov/geo/, accession number GEO: GSE196516). For quality control in single-cell RNA-seq data, we used the R package Seurat [58] to exclude cells with unique feature counts over 7500 or less than 200, as well as cells with a percentage of mitochondria genes greater than 25. We then employed the “LogNormalize” method with a scale factor of 10000. For ATAC-seq data, we used the Signac [59] package for quality control, during which we varied the number of peaks to exclude low quality cells. Note that we filtered out cells with transcriptional start site (TSS) enrichment score less than 1, or approximate ratio of mononucleosomal to nucleosome-free fragments greater than 1.2, or ratio reads in genomic blacklist regions greater than 0.006. The following table represents the number of cells before and after QC, as well as hyperparameter choices. Subsequently, we used scANANSE [13], which is an extension of ANANSE designed for single-cell sequencing data. We merged RNA and ATAC objects from two different days using days as clusters as input for scANANSE. Note that the ATAC objects were merged in without a common feature set because there are no common cells between days. After running scANANSE, we obtained the influence file as well as networks for each time point. As for expression data, after quality control of scRNA-seq expression data, we converted the expression level of each day into bulk data by taking the average expression of genes across all cells.

**Table A1:** Metric choices during ATAC-seq data preprocessing.

| Days | # cells before QC | # cells after QC | nCount_peaks |
| --- | --- | --- | --- |
| D2 | 7051 | 6986 | 9k-75k |
| D3 | 6755 | 6636 | 9k-75k |
| D4 | 5019 | 4706 | 9k-75k |
| D5 | 1651 | 1526 | 9k-200k |
| D7 | 669 | 615 | 9k-200k |
| D10 | 3116 | 2462 | 9k-100k |
| D15 | 5068 | 2020 | 9k-100k |

## Appendix B Details on parameter selection

**Fig. B1:**
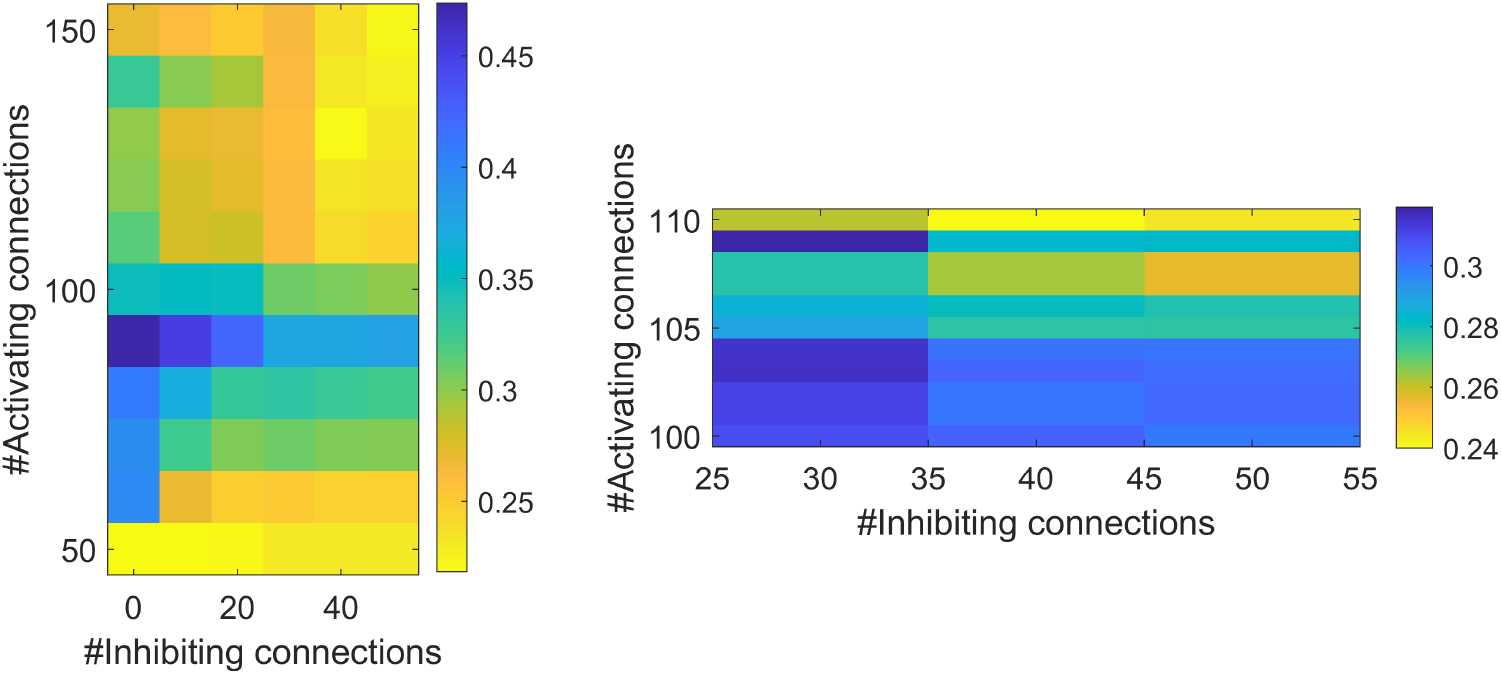
Average value of loss function over all nodes in chondrogenic differentiation model. **left** Value over large grid. **right** Value over zoomed in grid, showing cut-off value where fit improves. This cut-off is a key indicator for the choice of activating interactions when running DANSE.

**Fig. B2:**
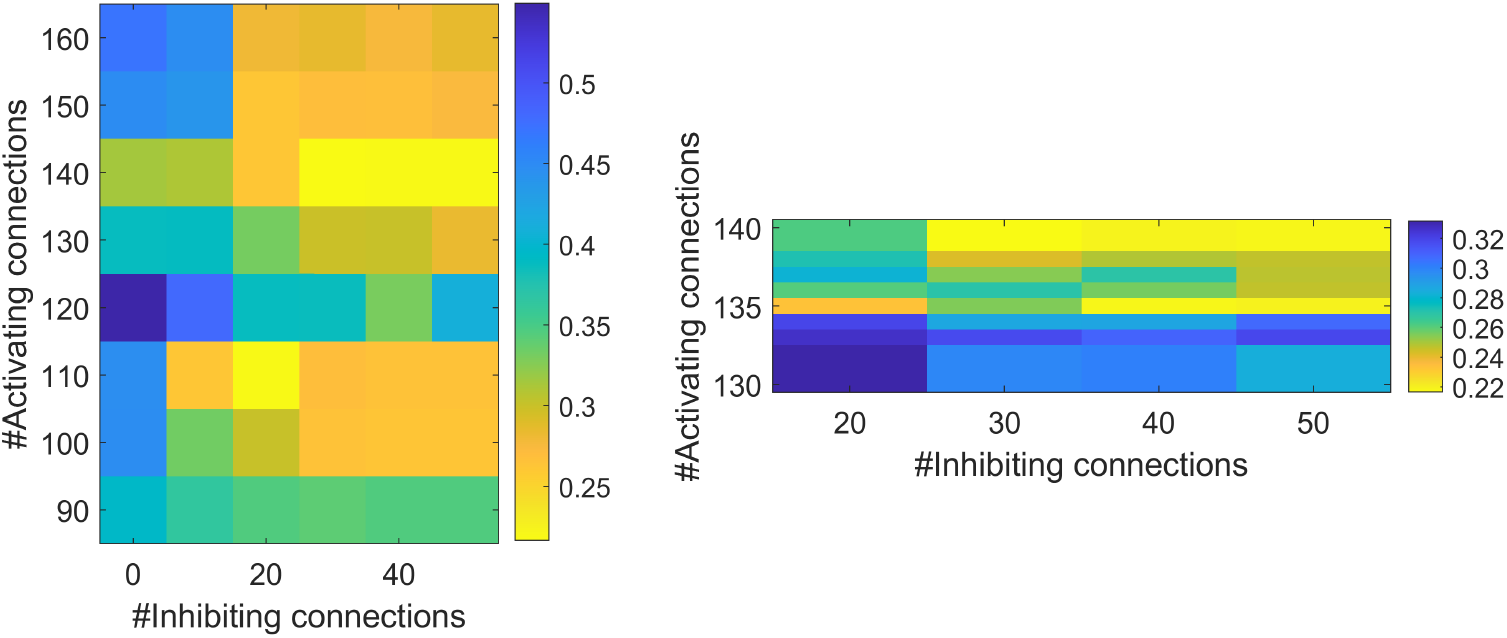
Average value of loss function over all nodes in epicardioid differentiation model.**left** Indication that model performs best with around 200 activating and 40 inhibiting connections. **right** Value over zoomed in grid, indicating 200 activating and 44 inhibiting connections to be the reasonable cut-off configuration

**Fig. B3:**
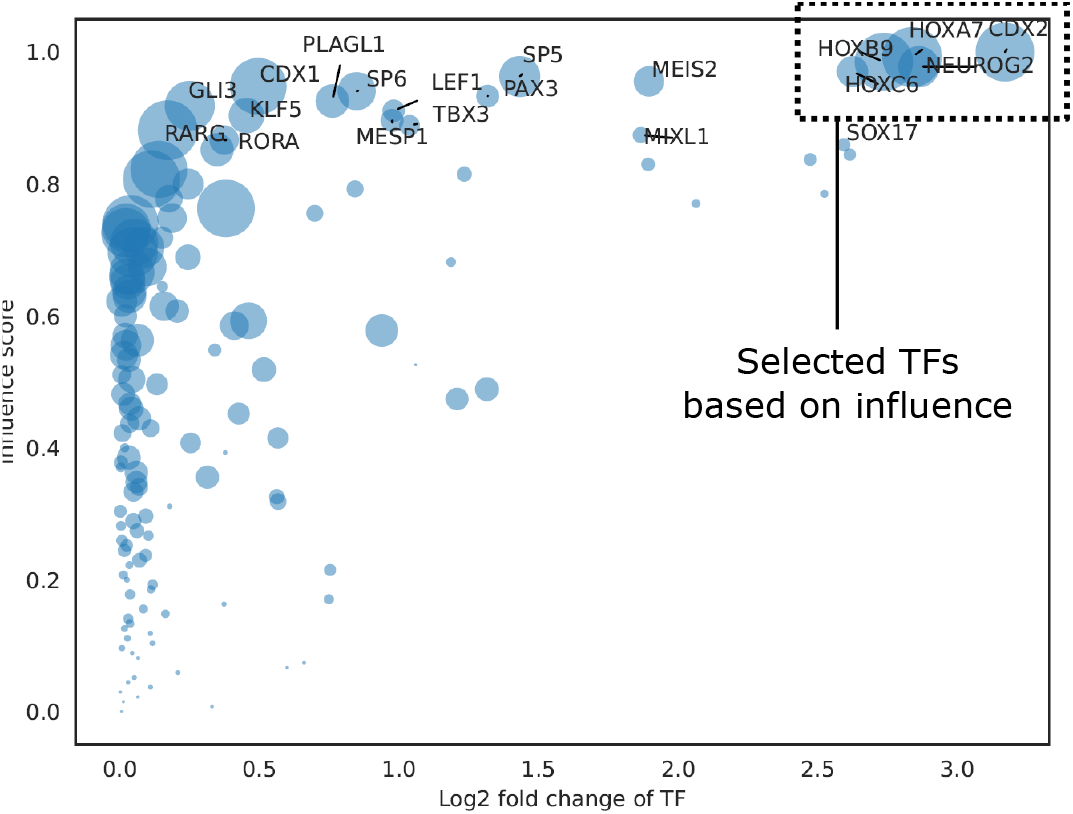
Influence plot for D0 → D2 in the The horizontal axis indicates fold change of expression level, whereas the vertical axis indicates change of the influence score, as determined using ANANSE [11]. The size of each dot indicates the expression level in the target cell population. The top 5 TFs (all selected TFs for the network from this influence plot) are indicated. The threshold of how many TFs are selected can be adjusted in DANSE, and desired TFs can be manually added.

**Fig. B4:**
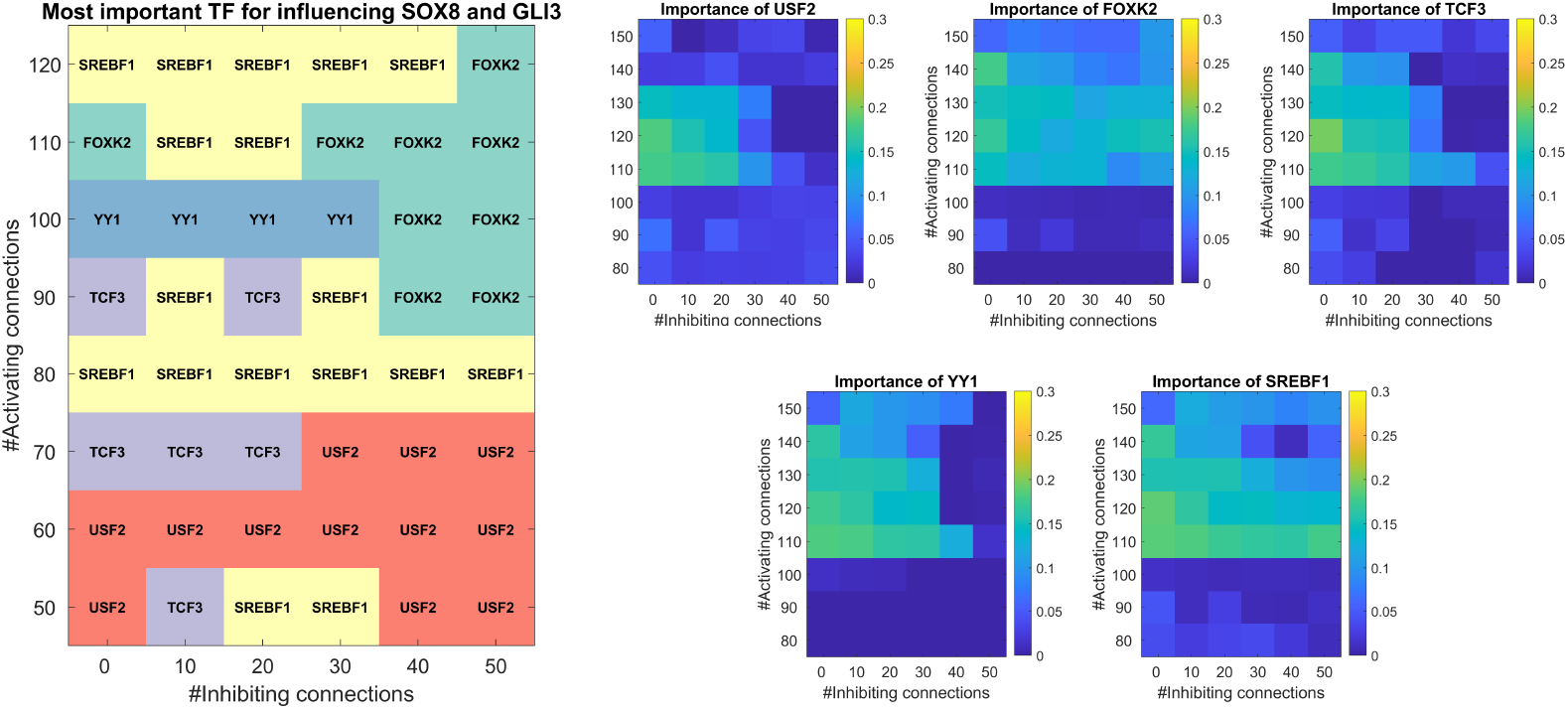
The most important TF as identified by the dynamical model constructed using DANSE for the chondrogenesis data [21] for different choices of activating and inhibiting connections. These TFs are classified as important by their influence on SOX8 and GLI3 expression in the constructed computational model. Here, only one fit is considered per grid point, which may cause minor differences in results compared to the statistics over 10 fits as shown in Figure 3. **left**. Overview of most important TF influencing SOX8 and GLI3 as identified for different choices for the number of activating and inhibiting connections. **right**. Importance for influencing SOX8 and GLI3 as calculated for different TFs. The calculation is shown for a varying number of selected activating and inhibiting connections.

**Fig. B5:**
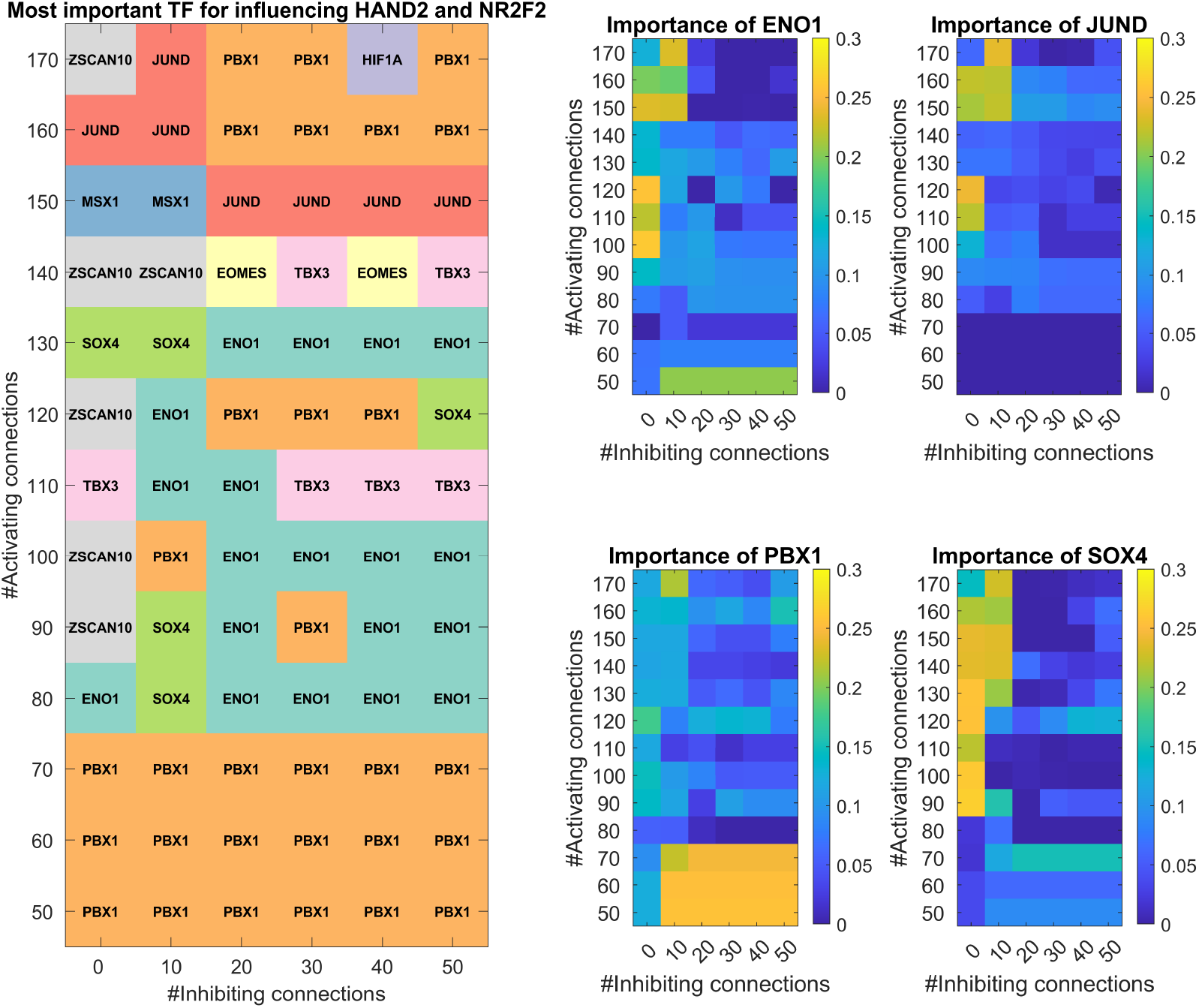
The most important TF as identified by the dynamical model constructed using DANSE for the epicardioid differentiation data [23] for different choices of activating and inhibiting connections. These TFs are classified as important by their influence on HAND2 and NR2F2 expression in the constructed computational model. Here, only one fit is considered per grid point, which may cause minor differences in results compared to the statistics over 10 fits as shown in Figure 4. **left**. Overview of most important TF influencing HAND2 and NR2F2 as identified for different choices for the number of activating and inhibiting connections. **right**. Importance for influencing HAND2 and NR2F2 as calculated for different TFs. The calculation is shown for a varying number of selected activating and inhibiting connections.

**Fig. B6:**
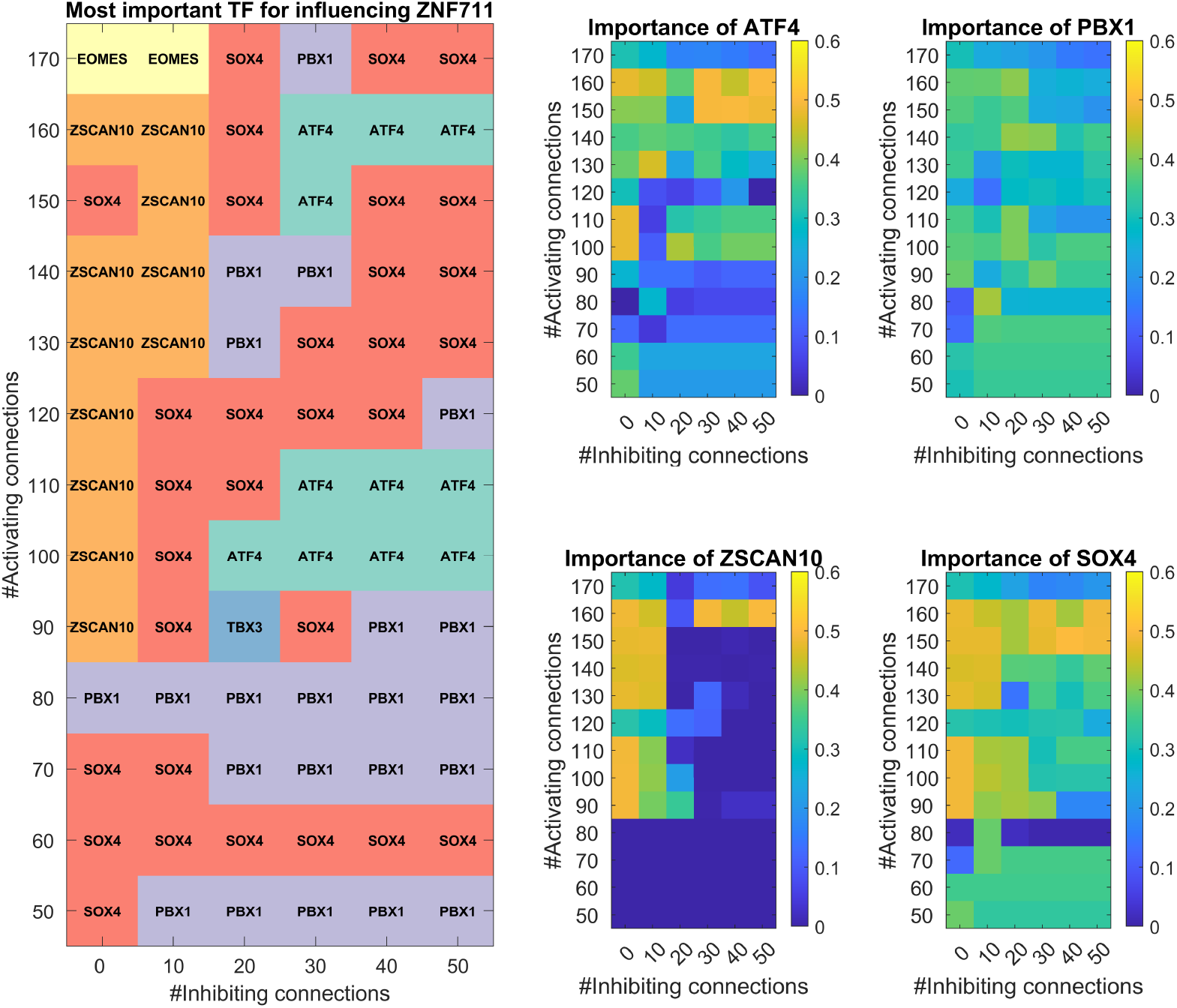
The most important TF as identified by the dynamical model constructed using DANSE for the epicardioid differentiation data [23] for different choices of activating and inhibiting connections. These TFs are classified as important by their influence on ZNF711 expression in the constructed computational model. Here, only one fit is considered per grid point, which may cause minor differences in results compared to the statistics over 10 fits as shown in Figure 4. **left**. Overview of most important TF influencing ZNF711 as identified for different choices for the number of activating and inhibiting connections. **right**. Importance for influencing ZNF711 as calculated for different TFs. The calculation is shown for a varying number of selected activating and inhibiting connections.

## Appendix C Full list of selected transcription factors

### C.1 The chondrogenesis model

**Fig. C7:**
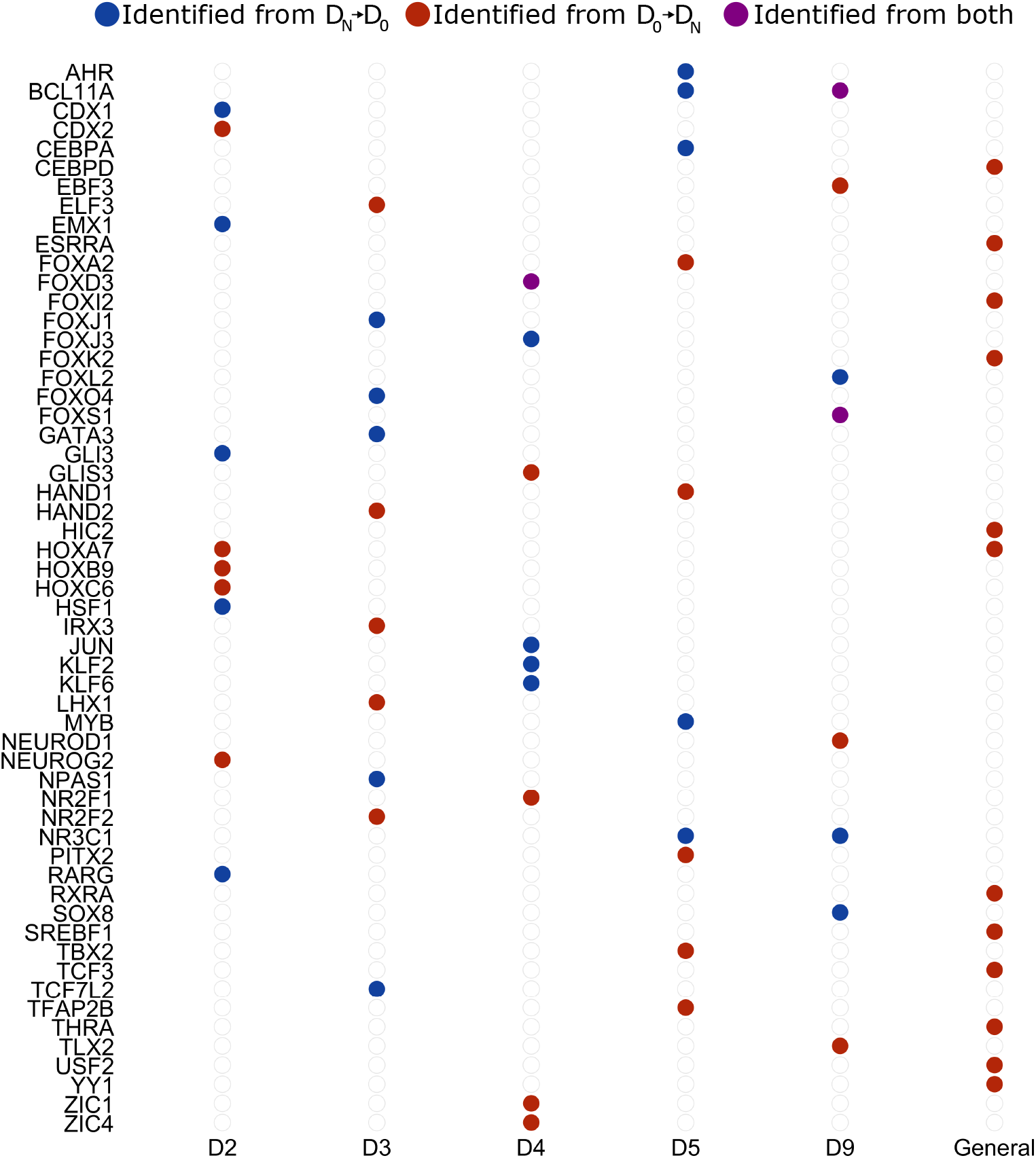
Overview of selected genes for the model based on the data from [21].

**Fig. C8:**
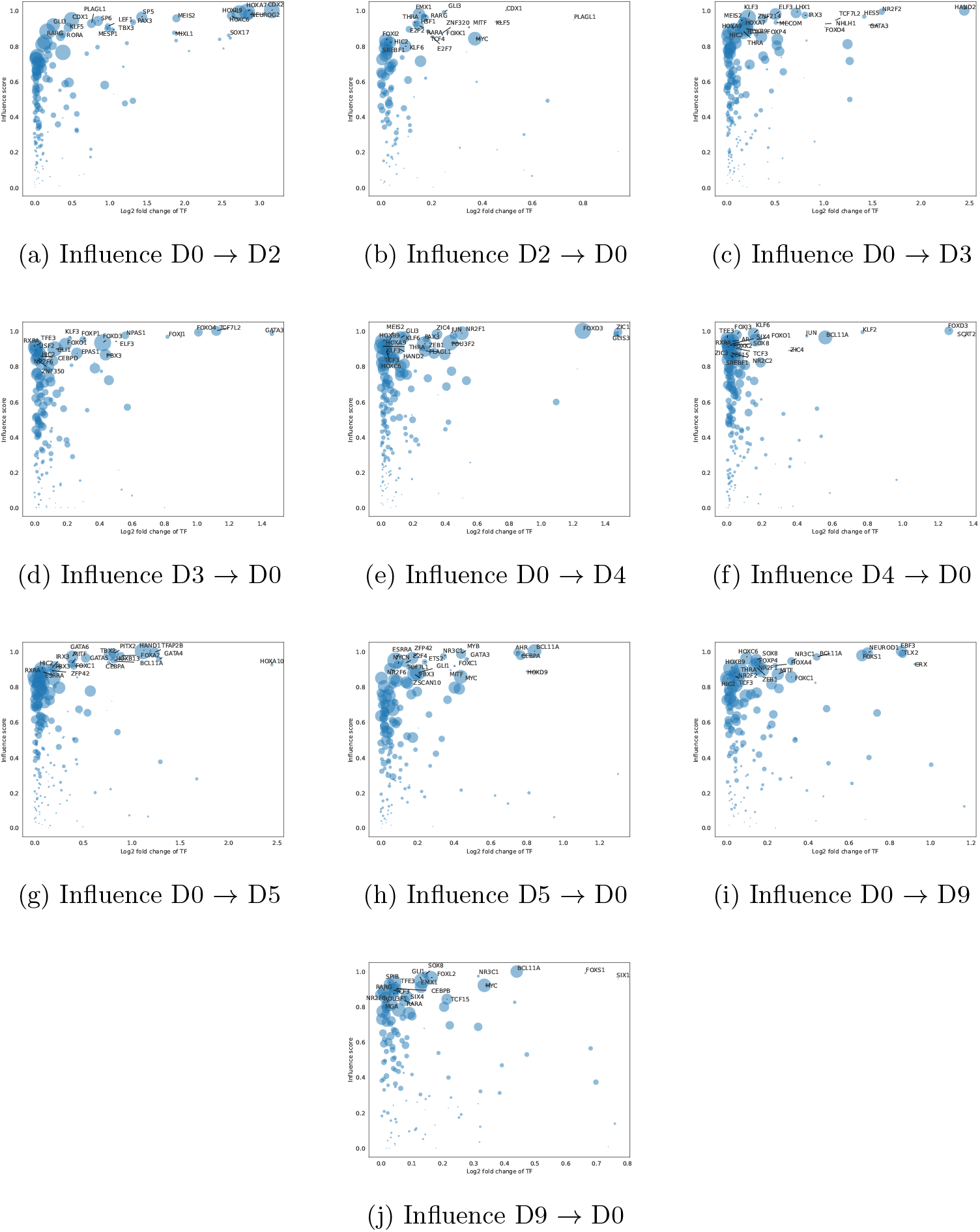
All influence plots for Kawata data. The horizontal axis indicates fold change of expression level, whereas the vertical axis indicates change of the influence score, as determined using ANANSE [11]. The size of each dot indicates the expression level in the target cell population.

### C.2 The cardiogenesis model

**Fig. C9:**
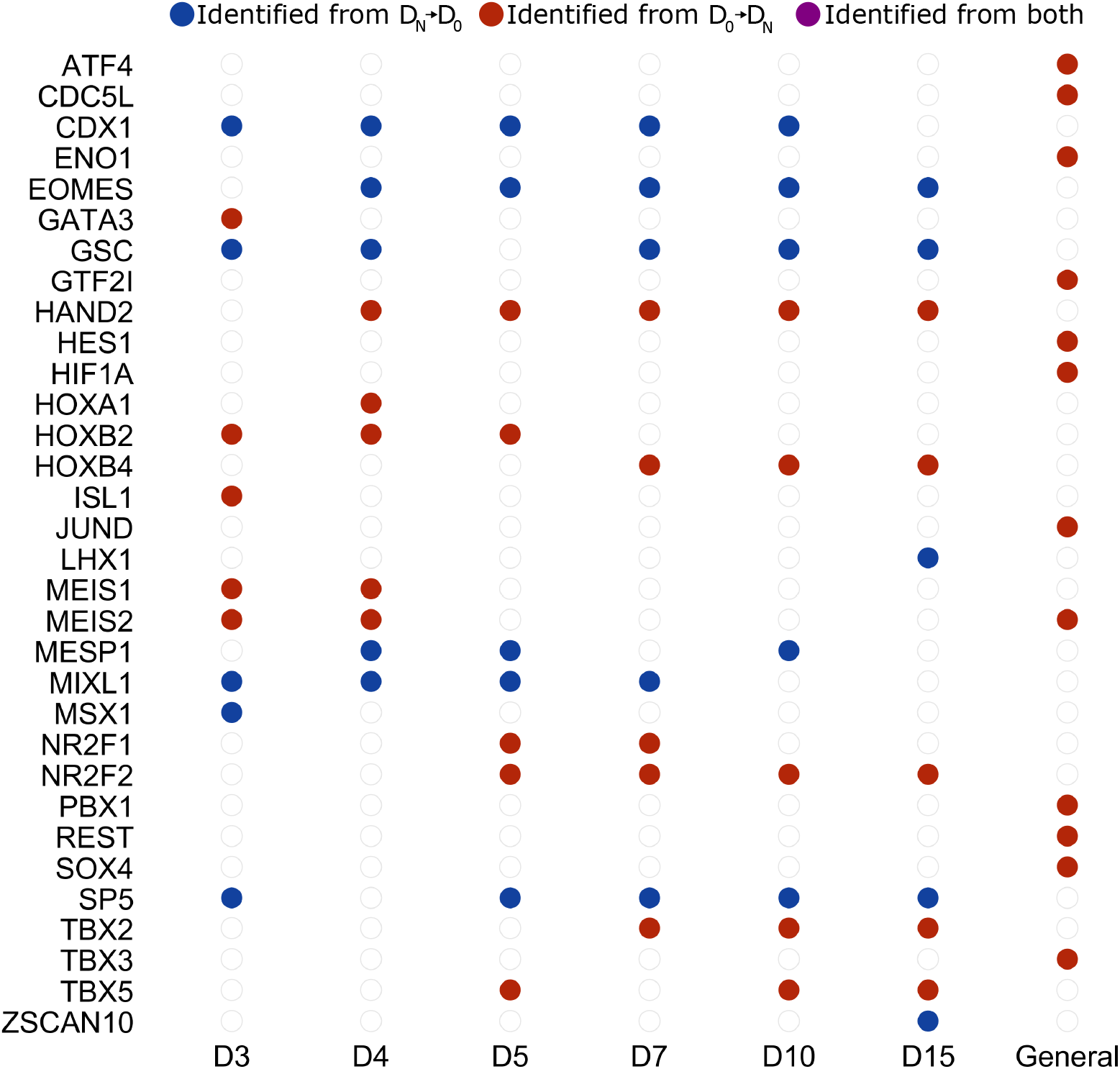
Overview of selected genes for the model based on the data from [23].

**Fig. C10:**
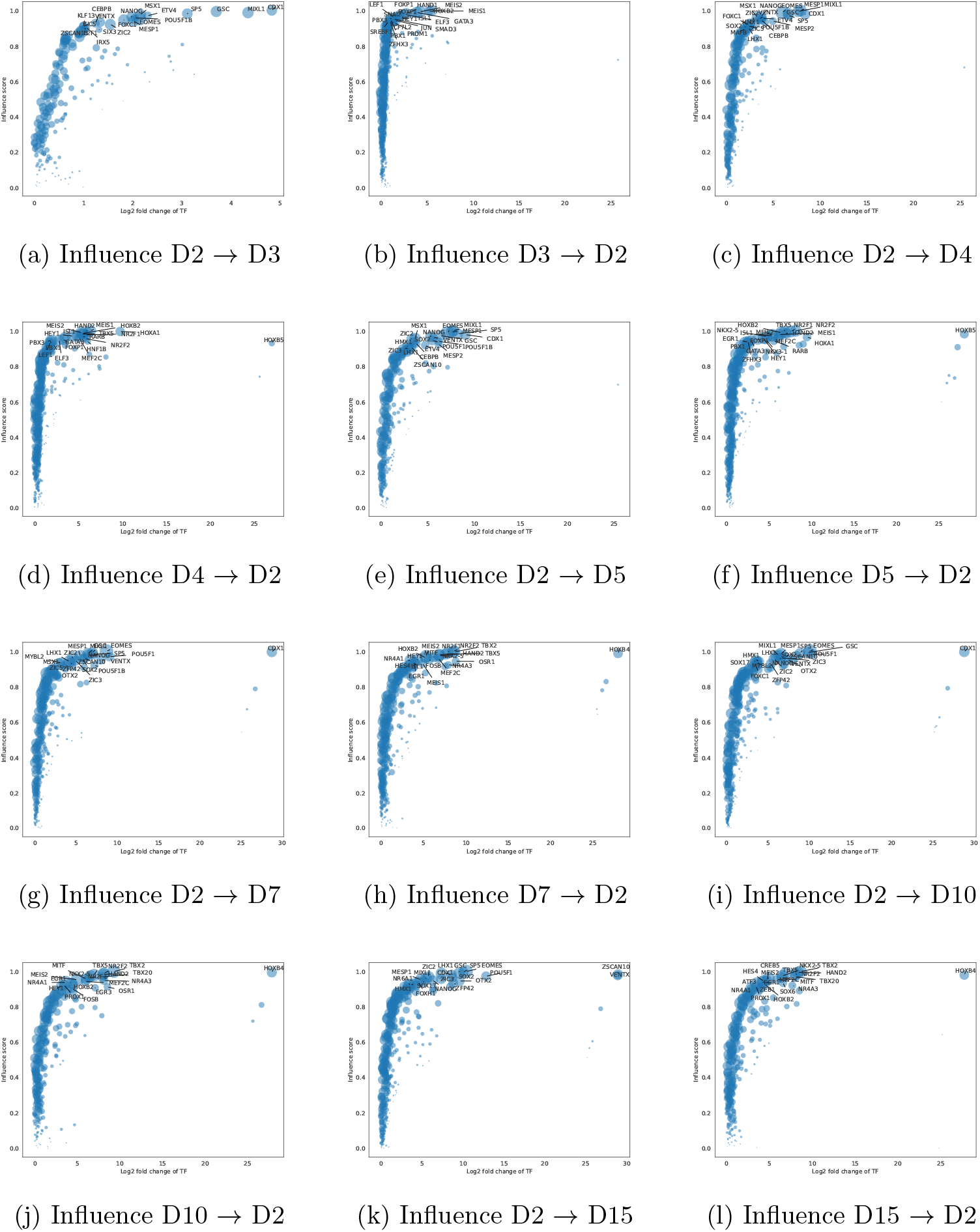
All influence plots for Meier data. The horizontal axis indicates fold change of expression level, whereas the vertical axis indicates change of the influence score, as determined using ANANSE [11]. The size of each dot indicates the expression level in the target cell population.

